# Evolving initial conditions: an alternative developmental route to morphological diversity

**DOI:** 10.64898/2026.04.01.715779

**Authors:** Shannon E. Taylor, James E. Hammond, Berta Verd

**Affiliations:** Department of Biology, University of Oxford

## Abstract

Phenotypic diversity is often thought to arise from the evolutionary modification of developmental processes. However, developmental processes are tightly coupled in space and time, with each process beginning from conditions set by the one before it. While we know from dynamical systems theory that initial conditions can significantly affect a system’s out-come, their importance as a source of phenotypic evolvability has been largely overlooked. Here we show for the first time, that phenotypic evolution can proceed through changes in developmental initial conditions while the underlying developmental process remains conserved. Somitogenesis is the process by which vertebral precursors, known as somites, are periodically patterned in the pre-somitic mesoderm (PSM). Somitic count (total number of somites) is thought to diversify through the evolution of components of somitogenesis such as the tempo of the segmentation clock or the mechanisms driving axial morphogenesis. Using two closely related species of Lake Malawi cichlid fishes that differ in vertebral counts, we show that somite count evolution has happened without changes to somitogenesis itself, but instead, by altering the size of the PSM at the onset of this process. This work will expand what we consider developmental drivers of phenotypic evolution and highlight the importance of comparative studies to understand the diversification of phenotypes.

## INTRODUCTION

One of the fundamental and most longstanding questions in biology is that of how phenotypes evolve to generate the remarkable diversity of forms that we see in the natural world. This ‘evolvability’ underlies life’s ability to adapt to ever-changing environments and exploit new ecological opportunities, resulting in a spectacular range of morphologies and innovations across the tree of life. The evolutionary modification of developmental processes has been found to often underlie the evolution of the adult phenotype, yet many of the mechanisms by which developmental change leads to phenotypic evolution remain unclear.

Development can evolve in many ways, and alterations to the timing or duration of developmental events, known as heterochronic shifts, are among the most well studied. Hete-rochronic shifts have been found to play a role in the evolution of a broad range of pheno-types such as beak shape across Darwin’s finches (Mallarino et al., 2012), bird skulls (Bhullar et al., 2012) and crocodilian snouts (Morris et al., 2019), all of which arose from changes in the timing of cranio-facial development. Similarly, evolutionary changes in developmental pattern formation - heterotypic shifts - also contribute toward generating phenotypic di-vergence as exemplified by the evolution of limblessness in snakes, which has been driven by spatial changes in gene expression during development (Cohn & Tickle, 1999).

Gene regulatory networks play a central role in establishing the temporal and spatial organisation of developmental patterning, and evolutionary changes in these networks can have profound phenotypic consequences. Heterochronic shifts caused by changes in biochemical network parameters such as protein degradation rates, have been seen to underlie inter-species differences in the tempo of neural differentiation (Rayon et al., 2020) and the frequency of the segmentation clock (Matsuda et al., 2020; Lázaro et al., 2023). Changes to the architecture of gene regulatory networks, including substantial ones such as gene co-option, underpin some of the most striking cases of morphological innovation, such as the evolution of beetle horns and butterfly eyespots (Hu et al., 2018; Murugesan et al., 2022). Recently, attention has turned to the possibility that the evolution of morphogenesis itself - independent of the underlying patterning mechanisms - may also contribute towards phenotypic diversification (Münster et al., 2019; Petridou et al., 2021; Busby & Steventon, 2021; Fulton et al., 2022a; Serrano Nájera & Weijer, 2023), although concrete empirical examples of this are still scarce.

It is well established that modifying the developmental processes patterning adult traits in the embryo often underlies the evolution of phenotypic diversity and novelty. However, how changes earlier in development might affect phenotypes patterned later remains poorly understood. Developmental processes are tightly coupled in space and time, with each process unfolding from the conditions set by the processes that preceded it. Theory, particularly dynamical systems theory (Jaeger & Monk, 2014; Verd et al., 2014; Crombach et al., 2016; Sáez et al., 2022; Kadiyala et al., 2025), has shown that initial conditions can exert a strong influence over system trajectories, significantly altering the final phenotypic state. Despite this, initial conditions have not yet been explicitly considered an evolvable parameter in developmental evolution in the same way that rates, duration or network architecture have. This gap limits our understanding of how variation arising earlier in development propagates to influence later-forming traits and ultimately contributes to morphological evolution.

Vertebral count is one of the most diverse traits across vertebrates, varying from less than 10 in some frogs to several hundred in snakes, all while being remarkably constant within any given species (Bucklow et al., 2025b; Hammond et al., 2025). Vertebrae develop from somites, paired blocks of mesodermal tissue formed in the early vertebrate embryo on either side of the developing spinal cord (Stickney et al., 2000; Holley, 2007; Oates et al., 2012). Somites are patterned periodically as the vertebrate embryo elongates posteriorly during a process known as somitogenesis (Schröter et al., 2008; Oates et al., 2012). Their formation is preceded by travelling waves of gene expression which sweep across the presomitic mesoderm (PSM) from posterior to anterior to determine the position of the next somite boundary in the anterior PSM (Holley, 2007; Oates et al., 2012). These waves emerge from a network of coupled genetic oscillators known as the segmentation clock, which drives oscillatory gene expression dynamics at the single cell level and controls the tempo of somite formation (Schröter et al., 2012; Oates et al., 2012). The position at which oscillations stop in the anterior PSM is given by the wavefront, first proposed by Cooke and Zeeman in 1976 (Cooke & Zeeman, 1976) as a positional threshold that has since been proposed to be molecularly implemented by gradients of signalling molecules including Wnt and FGF (Oates et al., 2012). The wavefront recedes, coupled to the elongating PSM (Aulehla & Pourquié, 2010).

While somitogenesis is conserved across the vertebrate tree, the underlying gene expression dynamics and morphogenetic mechanisms have been seen to vary widely across species. There is a remarkable degree of plasticity in the genetic components of the segmentation clock, and its period has been shown to differ between species (Krol et al., 2011; Hammond et al., 2025), with recent *in vitro* studies showing that clock periods differ up to fourfold across mammalian species (Lázaro et al., 2023). The morphogenetic mechanisms driving PSM elongation have also been found to diverge across species; patterns of cell rearrangements, motility and proliferation have all been found to be evolvable (Lawton et al., 2013; Bénazéraf et al., 2010; Steventon et al., 2016; Thomson et al., 2021; Mongera et al., 2018; Romanos et al., 2024). Furthermore, the total duration of somitogenesis has also been shown to differ between species (Hammond et al., 2025). Together these data illustrate that somitogenesis is a highly evolvable and plastic developmental process, which explains why it has been assumed that its evolution is responsible for the evolution of vertebral counts.

Most of what we know about somitogenesis comes from studies in traditional model organisms such as mouse, chick and zebrafish (Oates et al., 2012; Bénazéraf, 2019). While the study of these systems has been very successful at elucidating general principles and important species-specific differences in somitogenesis, the last common ancestor of these species lived more than 400 million years ago. Divergence of these species over such a long timescale will obscure the developmental origins of evolvability: with co-evolution of many developmental processes, it is impossible to tell which processes evolve quickest or most readily. Therefore, to study the evolvability of somitogenesis we focus on a closely related species group, the Lake Malawi Cichlid fishes.

Lake Malawi cichlids are a powerful emerging model system in evolutionary developmental biology (Turner, 2007; Santos et al., 2023). Around 850 cichlid species have evolved in Lake Malawi from a single common ancestor in less than a million years (Malinsky et al., 2018). In this short amount of time, cichlids have colonised every available niche in the lake, exhibiting an astonishing degree of phenotypic diversity in traits such as body shape and size, colour and pattern, cranio-facial morphology, social and breeding behaviours, and vertebral counts which range between 28 and 40 (Bucklow et al., 2025a, 2025b; Santos et al., 2023). In this work, we take advantage of the close relatedness and spectacular diversity of Lake Malawi cichlids to investigate the developmental basis of vertebral count evolution. To do this, we compare somitogenesis in *Astatotilapia calliptera*, the putative ancestor of the Malawian cichlid radiation (Malinsky et al., 2018) which make 32 somites with *Rham-phochromis* sp. ‘chilingali’, one of the most elongated species in the lake catchment, which make 38 somites (Marconi et al., 2023), resulting in a 25% difference in somitic counts be-tween the two.

Our results show that somitic counts have evolved without evolving somitogenesis: we find that the tempo of somitogenesis is conserved, as are the mechanisms underlying PSM mor-phogenesis. Instead, we see that the initial conditions of the process have evolved dramatically, with the PSM being much larger at the onset of somitogenesis in *R*. sp. ‘chilingali’ than in *A. calliptera*. We see that this difference in the initial size of the tissue is enough to increase the total duration of the process, increasing the total number of somites patterned as a result without the need to evolve somitogenesis itself. Thanks to the short evolutionary distances separating our model species, we identify a case of evolution via the initial conditions of a developmental process, as opposed to the evolution of the process itself. We propose that this is a generally applicable mechanism of phenotypic evolution, particularly over short evolutionary timescales, one that may drive early diversification events during speciation.

## RESULTS

### The duration of somitogenesis differs between *A. calliptera* and *R*. sp. ‘chilingali’, but segmentation clock tempo is conserved

The simplest explanation for the increased number of somites (referred to also as somite numbers or somite/somitic counts) in *R*. sp. ‘chilingali’ would be an increase in either the rate or the duration of somitogenesis, or both (Gomez et al., 2008; Hammond et al., 2025). To investigate this, we quantified the total number of somites in embryos of known ages, staged in hours post fertilization (hpf) (Fig.1.A-M). We found that somitogenesis does not begin at the same time in both species: the first somite in *A. calliptera* appears at 33hpf, while in *R*. sp. ‘chilingali’ it appears five hours later, at 38hpf. This indicates that *R*. sp. ‘chilingali’ embryos take longer to complete the blastulation and gastrulation phases of development than those of *A. calliptera*. Additionally, we see that somitogenesis takes longer to complete in *R*. sp. ‘chilingali’ than in *A. calliptera*: Fig.1.N shows that *A. calliptera* embryos complete somitogenesis at 68hpf (i.e. possess 33 somites), while *R*. sp. ‘chilingali’ will not have the complete set of 39 somites until 80hpf. The duration of somitogenesis is therefore increased in *R*. sp. ‘chilingali’ with respect to *A. calliptera*, taking 42 hours compared to 35, an extra seven hours in *R*. sp. ‘chilingali’ than in *A. calliptera*.

**Figure 1:**
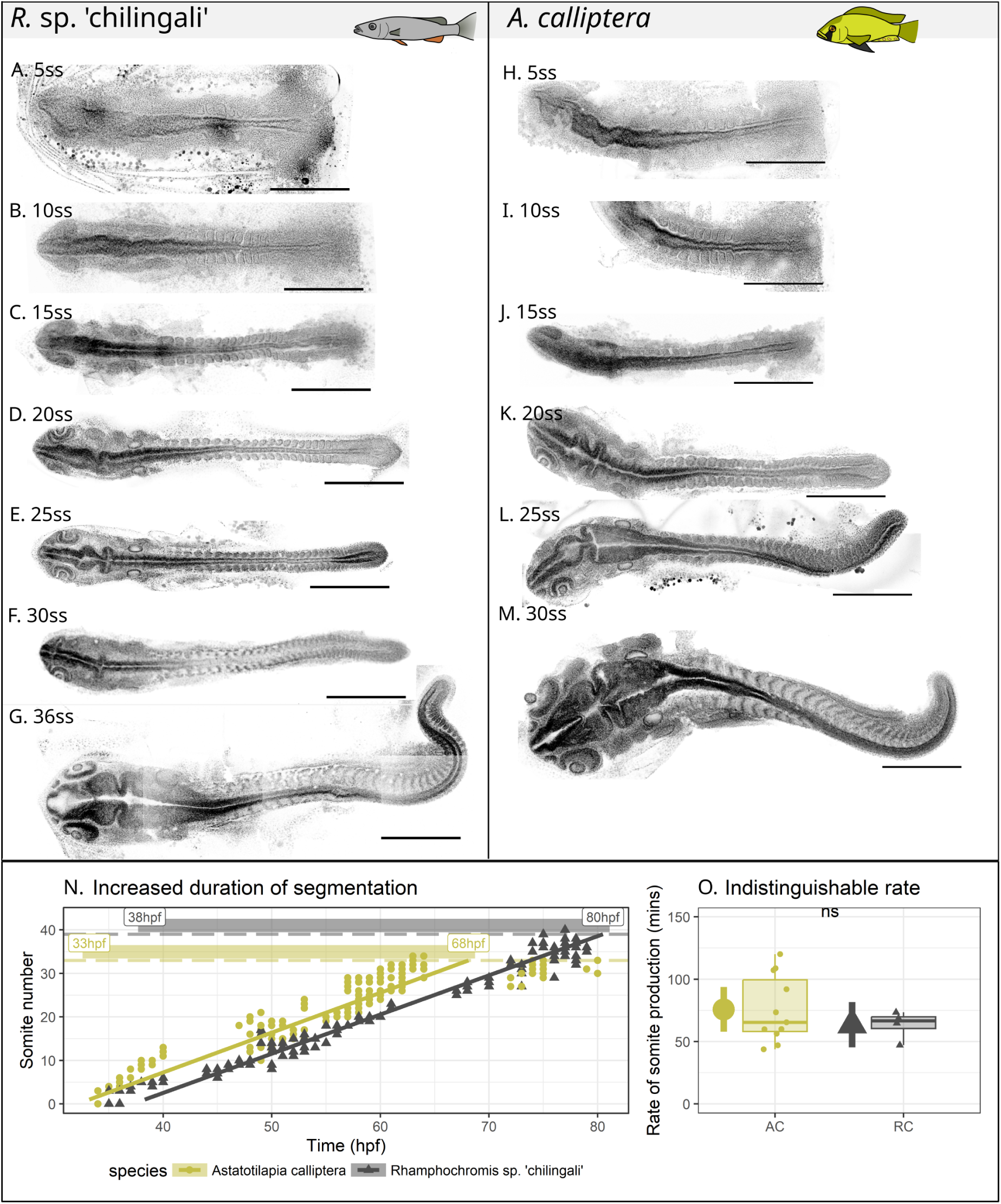
Overview of cichlid somitogenesis. **A-M:** single optical section of flat-mounted of DAPI-stained (black) cichlid embryos at different somite stages (ss). All embryos are a dorsal view, oriented anterior left, to the same scale. Scale bar: 500 m. **N**. Number of somites versus time in hours post fertilisation (hpf) in both species. The linear regression was constructed for both species using ordinary least squares and the best fit equation is displayed at the bottom right. Yellow: *A. calliptera*, black: *R*. sp. ‘chilingali’. **O**. Somitogenesis rate in both species, computed for each clutch with more than two measurements/embryos available. *A. calliptera* embryos older than 70hpf are excluded from analysis, as they have finished producing somites, as are clutches with no limited temporal resolution (less than two hours, and less than three samples).

Interestingly, however, while the total duration of somitogenesis is increased in *R*. sp. ‘chilingali’ with respect to *A. calliptera*, it appears that the rate of somitogenesis does not differ between these two species. In *A. calliptera* a somite forms every 65 minutes on average, and in *R*. sp. ‘chilingali’ a somite forms every 67 minutes; when comparing across different clutches this difference was not found to be statistically significant (p=0.42, two-sided t-test, Fig.1.O., Fig.S1.).

Our data show that *R*. sp. ‘chilingali’ has evolved increased somite counts by increasing the duration of somitogenesis instead of altering the rate of somite production, results that agree with prior work by Marconi and colleagues (Marconi et al., 2023), who show limited differences in the rate of somite production between these species.

### The speed of the wavefront and the rate of elongation are predicted to be conserved across species

We then wanted to establish whether the velocity *v* of the wavefront of somite determination, which we use as a proxy for the speed of axial elongation, differs between species. According to the clock and wavefront model (Cooke & Zeeman, 1976; Oates et al., 2012), this relationship should be captured by the formula 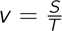 where *S* is the nascent somite length, and *T* is the tempo of somitogenesis . We measured the length of the nascent somite directly, showing that this is very similar between the two species (Fig.2A). Nascent somites in *A. calliptera* are consistently slightly longer at the same stage than those of *R*. sp. ‘chilingali’, with the *A. calliptera* somite being on average 2.7*µ*m longer (Fig.2A). However the effect size of this difference is small (p = 0.01; *η*^2^ = 0.05), indicating that this difference likely has limited significance to the embryo or rate of elongation. The volume of the nascent somite is statistically indistinguishable between species (p = 0.82, two way ANOVA also accounting for somite stage), and decreases over developmental time for both (Figure 2C, p < 0.001; *η*^2^=0.32).

**Figure 2:**
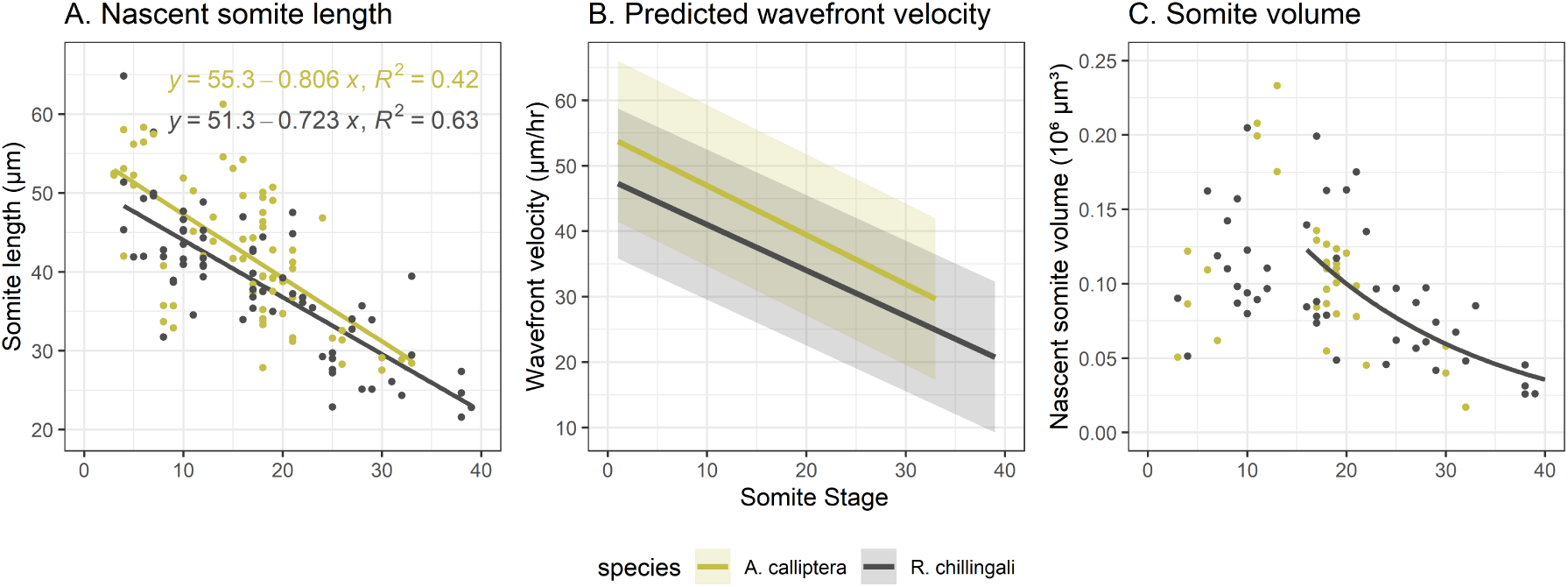
Somite size and wavefront velocity. **A**. *A. calliptera* have longer nascent somites than *R*. sp. ‘chilingali’. Somite antero-posterior length, measured from high-resolution confocal images of the PSM, is plotted against somite stage for both species. **B**. Predicted velocity of the wavefront is similar between species. **C**. Volume of the nascent somite is the same between species.

As expected, given that neither somite length nor segmentation rate differ markedly between species, the predicted velocity of the wavefront is also predicted to be very similar between the two, suggesting that the speed of axial elongation during somitogenesis might also be conserved between these two species (Fig.2.B).

### PSM size differs significantly at the onset of somitogenesis between *R*. sp. ‘chilingali’ and *A. calliptera*

We have shown so far that *A. calliptera* and *R*. sp. ‘chilingali’ differ in the timing of the onset of somitogenesis as well as in the total duration of this process (Fig.1.N&O), but that there is no evidence that evolution of segmentation clock or wavefront dynamics might have contributed towards the evolvability of somite number in Lake Malawi cichlids. Instead, we found that the duration of somitogenesis differs significantly between these species (Fig.1), driving the observed divergence in somite number. To understand this further, we characterized body axis morphogenesis during somitogenesis, particularly focusing on the morphogenesis of the pre-somitic mesoderm (PSM), the tissue that somites are produced from.

The anatomy of the PSM and the tailbud is similar between the two cichlid species, the only visually remarkable difference lies in the size of the tailbud at early stages of somitogenesis which already by eye appears to be much larger in *R*. sp. ‘chilingali’ with respect to *A. calliptera* (Fig.3A-F). We quantified the morphodynamics of the PSM, measuring both PSM volume and the number of cells in the PSM over the course of somitogenesis. We observe that the growth dynamics of this tissue change over time. Prior to 15 somites stage (ss), there is no statistical evidence that the PSM changes in volume over time in either species (p = 0.54; two way ANOVA testing the interaction between PSM volume and somite count alongside species). After 15ss, the data are well-approximated by an exponential decay curve in both species (Fig.3.C). It is probable that prior to 15ss, loss of tissue to the somites is balanced by lateral ingression of cells into the PSM, resulting in a PSM that does not change in volume, while the paraxial mesoderm, comprising already-formed somites as well as the PSM, is undergoing volumetric growth. This is different to other species surveyed (Lamprey, dogfish, zebrafish, mouse, and multiple species of snake) where the PSM is either increasing or decreasing in volume at any one time (Gomez et al., 2008; Steventon et al., 2016).

**Figure 3:**
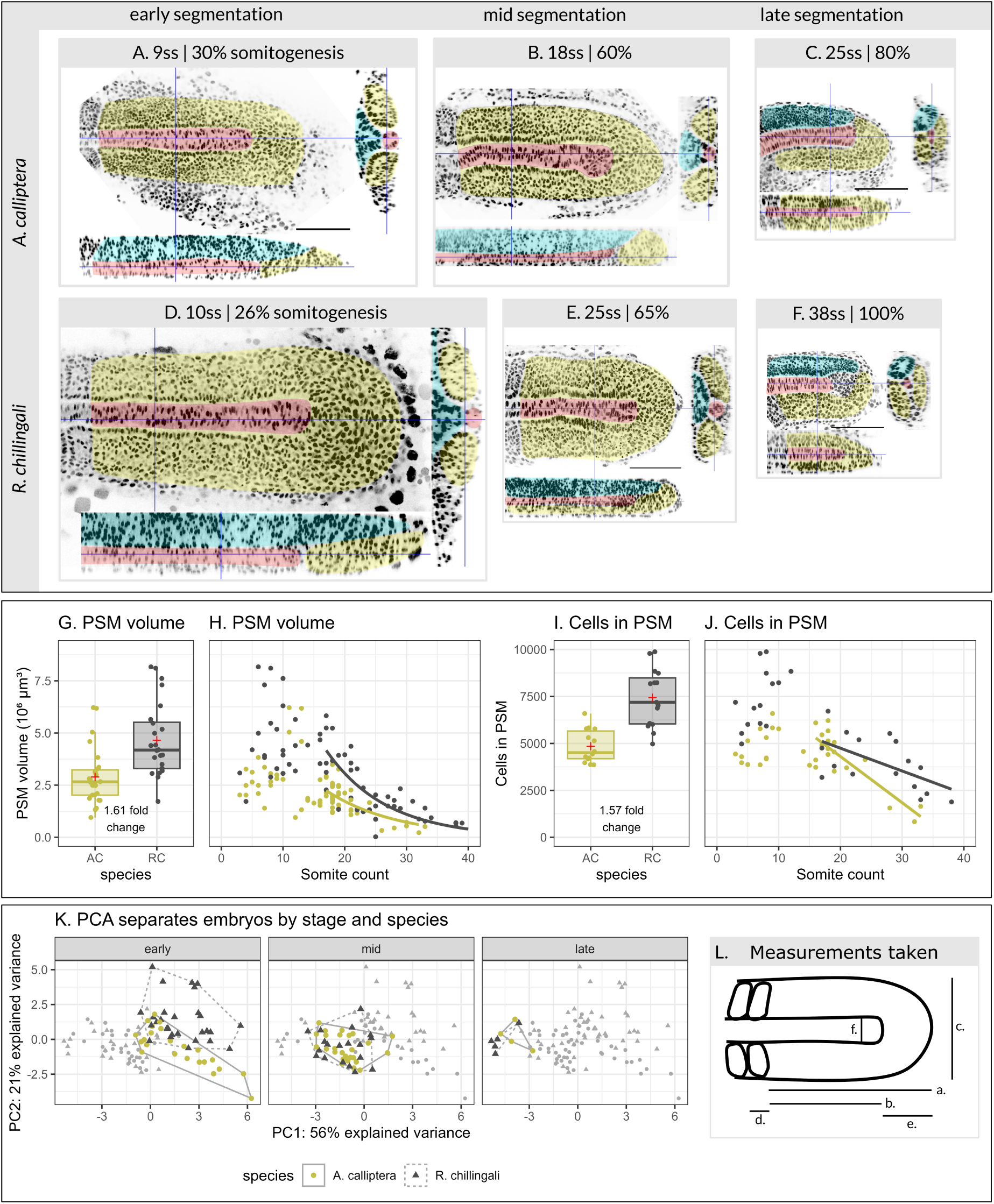
PSM size differs between species only at the onset of segmentation. **A-F.** Single confocal slices showing the anatomy of the PSM over time in *A. calliptera* and *R*. sp. ‘chilingali’. In each panel, the middle image depicts a sagittal slice of the PSM at the level of the notochord; the bottom image depicts a dorsoventral slice of the PSM through the notochord, from neural tube to notochord, and the right image depicts a dorsoventral slice of the PSM oriented medio-laterally, at the middle of the PSM. Tissues are false-coloured as follows. PSM: yellow. Notochord: red. Neural tube: cyan. DAPI signal is shown in greyscale. Embryos are oriented anterior to the left; scale bar 100m. All embryos to the same scale. Caption continued on following page. **G**. Quantification of PSM volume for both species prior to 15ss. Box-plots represent the upper and lower quartiles of the data, with the solid line representing the median of the data. Red cross represents the mean of the data. Whiskers represent the largest value no further than 1.5 times the inter-quartile range of the plot. **H**. PSM volume over time. PSM volume was measured manually as described in the Methods section. The trend line was fit to an exponential decay function using nonlinear least squares. **I**. Quantification of number of cells in the PSM for both species prior to 15ss. Boxplots are laid out as for G. Cells were counted using a custom automatic script defined in the Methods. **J**. Number of cells the PSM over time. The trend line was fit to a linear model using ordinary least squares. **K**. Principal component analysis (PCA) of given variables separates species only at early stages of somitogenesis (prior to 15ss). **L**. Schematic of variables used for PCA. a: PSM length. b: Notochord length. c: PSM width. d: nascent somite length. e: Mesoderm progenitor zone (MPZ) length. f: notochord length. Also measured and included were PSM depth, PSM length to width ratio, PSM length to depth ratio, PSM width to depth ratio, and PSM length to somite length ratio. All data points are plotted in grey, and data points for the indicated stage are coloured: (*A. calliptera* yellow, *R*. sp. ‘chilingali’ grey).

*A. calliptera* and *R*. sp. ‘chilingali’ differ in the size of the PSM at early stages of somitogenesis, particularly with regards to volume. Prior to 15ss, the volume of the *R*. sp. ‘chilingali’ PSM is on average 61% larger than that of *A. calliptera* (Fig.3.E; p < 0.001, *η*^2^ = 0.25, two way ANOVA). After 15ss, the exponential decay trend lines describing both species’ development show that the volume of the *R*. sp. ‘chilingali’ PSM remains always larger than that of *A. calliptera*. The number of cells in the PSM also differ at early somitogenesis be-tween both species: *R*. sp. ‘chilingali’ have 57% more cells than *A. calliptera* prior to 15ss (p <0.0001, *η*^2^ = 0.63, Fig.3.J). The number of cells in the PSM is also always consistently larger in *R*. sp. ‘chilingali’ after 15ss (p = 0.047; *η*^2^ = 0.12, Fig.3.J).

In addition, we also measured the length of the whole embryo, the length of the PSM, and the proportions of the embryo over developmental time. We found that *R*. sp. ‘chilingali’ are consistently larger than *A. calliptera*, with larger embryonic and PSM lengths (Fig.S2.A-D). While the head occupies a similar proportion of the embryo between species (FigS2E,G), the PSM occupies a larger proportion of the embryonic length in *R*. sp. ‘chilingali’ than in *A. calliptera*. Thus, the *R*. sp. ‘chilingali’ PSM is disproportionately larger than that of *A. calliptera*.

We performed a detailed quantitative comparison of the shape of the PSM over the course of somitogenesis in both species by measuring the dimensions of the different regions in the cichlid unsegmented tailbud: we measured the PSM length, width and depth, the length of the notocord, the length of the putative mesodermal progenitor zone (MPZ), the nascent somite size, and also the ratios between these values (11 metrics in total, Fig.3.L).

Principle component analysis (PCA) on this dataset separates embryos by stage, with early stage embryos occupying the middle to middle right of the graph plotting PC1 and PC2, while midsomitogenesis embryos are placed in the middle and late somitogenesis embryos lie to the left (Fig.3.K). Importantly, *A. calliptera* and *R*. sp. ‘chilingali’ cluster together at mid and late stages of somitogenesis, only ever segregating by species at early stages of somitogenesis (Fig.3.K) with *R*. sp. ‘chilingali’ occupying the top right section of the plot (high in PC2) and *A. calliptera*, the bottom right of the plot (low in PC2). This suggests that the morphometric differences in tailbud shape between species are most pronounced and only significant at early stages of somitogenesis.

We next sought to uncover whether species differ in specific dimensions of the PSM. We find that within species, all tailbud measurements are significantly associated with somite stage (Table S1). After correcting for multiple testing and accounting for somite stage, only PSM length, notocord length, PSM length to somite length ratio, and MPZ length differ be-tween species (Fig.S3, Tables S1, S2). All of these are ultimately a consequence of the larger PSM length in *R*. sp. ‘chilingali’: notochord length is tightly correlated with PSM length, and the difference in PSM length between species with the conserved somite lengths explains the different PSM length to nascent somite length ratios. Finally, the increased size of the MPZ may be due to a larger pool of progenitor cells in *R*. sp. ‘chilingali’.

We next used these data to explore whether species differences are established at the onset of segmentation, or instead involve changes to the process of somitogenesis. For example, if species differ in PSM length, is this difference present at the onset of somitogenesis which would be reflected if the intercept of the slopes of the growth curves differed, or does the rate of change of PSM length differ between species, potentially indicating that the mechanism underlying PSM length changes could differ between species. For all the measurements which differ between species, our data supports the first possibility: the intercept differs, but not the rate of change over the course of development. This indicates that the morphogenetic processes are conserved but that the initial conditions have evolved (Fig.S3, Tables S1, S2).

Taken together, these analyses show that these two cichlid species differ most markedly at the earliest stages of somitogenesis. Even traits, such as MPZ size and PSM length, that do differ between species, differ most in their initial values and not in their rate of change over the course of somitogenesis. This suggests that while the initial conditions of this developmental process differ between species, the processes driving axial elongation during somitogenesis appear to be conserved in both cichlid species. Therefore, our data support a scenario where *R*. sp. ‘chilingali’ have evolved an increased number of somites by evolving initial PSM size at the onset of somitogenesis, rather than by altering the processes of PSM morphogenesis and axial elongation during somitogenesis.

### Both species begin somitogenesis with one somite, and add somites sequentially thereafter

In order to establish that total somitic counts indeed depend on the size of the PSM at the onset of somitogenesis, it was important to establish whether either species forms significant numbers of segments simultaneously at the onset of somitogenesis. We fixed embryos at hourly intervals at early stages of somitogenesis. When obtaining embryos with 0-4 somites for both species, these clearly show that neither species forms any somites simultaneously, consistent with findings in zebrafish and chick (Fig.S4, Fig.S5, Fig.4) (Schröter et al., 2008; Maia-Fernandes et al., 2024). Furthermore, we found no difference in the early versus later rates of segmentation nor between species: while the breeding behavior of cichlids means that development within a clutch is asynchronous and that there is variation in the number of somites represented at each time-point, it appears that somites also appear approximately once every hour at the onset of segmentation, as they do at later stages (Fig.S4 and Fig.S5).

**Figure 4:**
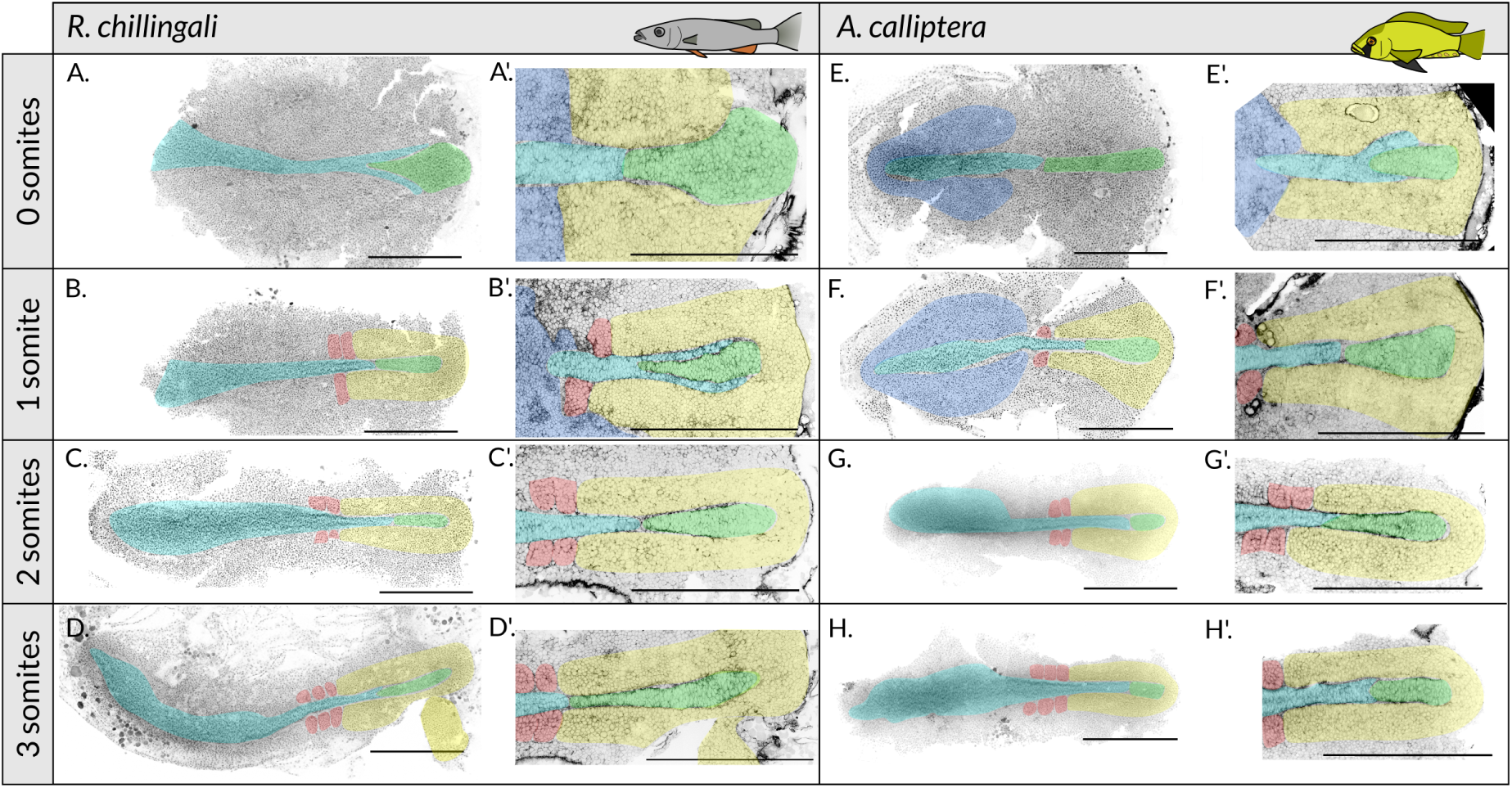
Anatomy of the early embryo and unsegmented region at the onset of segmentation. **A-H.** Whole-mount, dissected cichlid embryos stained with DAPI (nuclei) and phalloidin (actin). Images to the left (A-H) are shown complete, with nuclei in grey. Images to the right, **A’-H’**. magnify the unsegmented region, and show actin/phalloidin in grey. Individual tissues are false-coloured as follows: neural tube: cyan; notocord/notocord projenitors: green; somites: red; unsegmented mesoderm: yellow; ectoderm: dark blue. Scale bar: 500*µm*. All embryos are oriented with anterior to the left and are shown in a dorsal view.

Despite the fact that *R*. sp. ‘chilingali’ embryos at the onset of somitogenesis (0-3ss) are older (in hpf) than their *A. calliptera* counterparts, the early anatomy of these embryos is indistinguishable between both species. Compared to embryos at later stages of somitogenesis, in early somitogenesis the PSM appears morphologically continuous with the lateral plate mesoderm, and somites have not yet epithelialised laterally (compare Fig.4 and Fig.3.A-F). The only obvious difference between species lies in the overall size of the PSM, where as already shown above, the PSM in *R*. sp. ‘chilingali’ is bigger than in *A. calliptera*. These data further support the notion that it is the initial size of the PSM that determines total somitic counts, and that species-specific differences in PSM size are established prior to the onset of somitogenesis.

### The molecular anatomy of the tailbud is indistinguishable between species

To test whether the observed similarities in PSM morphodynamics were reflected at the molecular level we characterized the expression of genes known to be important for PSM morphogenesis and somite patterning across vertebrates, in both *A. calliptera* and *R*. sp. ‘chilingali’. These included the gene *tbxta*, known as *T-Bra* in mouse, which is required for the specification both of the notochord and the posterior body in vertebrates (Schulte-Merker et al., 1992, 1994; Martin & Kimelman, 2008, 2010). As in other vertebrate embryos studied, *tbxta* is expressed in the notochord both in *A. calliptera* and *R*. sp. ‘chilingali’, as well as in the germ ring in early embryos (Fig.5A’, B’). At mid stages of somitogenesis, *tbxta* is expressed in the notochord and in the posterior-most region of the PSM. *tbxta* expression is similar between species, and indistinguishable when quantified at mid segmentation (Fig.5A’, B’, D’, E’, F).

**Figure 5:**
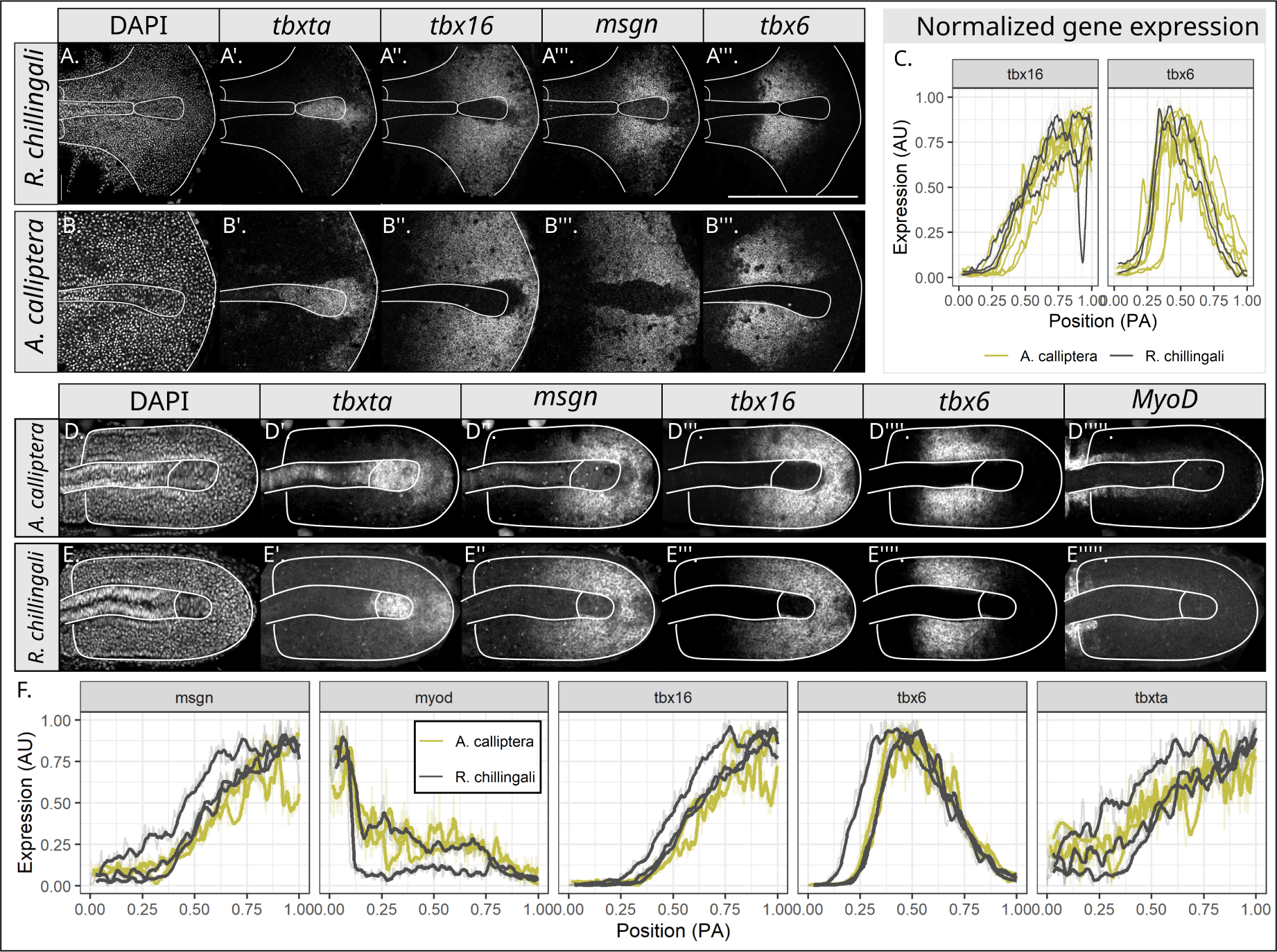
Patterning of the PSM is indistinguishable between species. **A-B.** Single confocal sections of very early somitogenesis stage embryos (3ss) simultaneously stained for DAPI, *tbxta*, *tbx16*, *msgn*, and *tbx6*. Scale bar: 500um. Thick white lines outline the nascent somites, notocord, neural tube, and unsegmented region boundary. **C**. Normalized gene expression measured across multiple embryos. Length and amplitude normalisation performed as described in the Methods section. In brief, *tbx6* and *tbx16* expression is measured along the middle of the PSM and across the same line for each gene. *N_AC_* : 6. *N_RC_* : 2. **D-E**. Single confocal sections of mid-segmentation stage embryos stained simultaneously for DAPI, *tbxta*, *tbx16*, *msgn*, *tbx6*, and *MyoD*. Thick white outlines outline the PSM, notocord, and neural tube. **F**. Quantifications of spatial gene expression for each gene at mid somitogenesis stages, normalised as described in the Methods section. The spatial expression of *tbxta*, *tbx16*, *msgn*, and *tbx6* was measured along the same line, across the midline of the embryo. The expression of *MyoD* is measured along the adaxial layer of cells. *N_AC_* = 2. *N_RC_* : 3. All embryos are oriented anterior left and shown in a dorsal view. Different channels of the same embryo are shown in each row, except for B”’.

The genes *tbx16* and *msgn* are required for correct cell motility patterns in the zebrafish tailbud, where they are expressed in a posterior-to-anterior gradient (Manning & Kimelman, 2015). This expression pattern is conserved in cichlids, and when quantified, the slope and position of the expresssion domain is indistinguishable between both species at early and mid-segmentation (Fig.5A”’, A”, B”, B”’, C, D”, D”’, E”, E”’, F). The gene *tbx6* has been found to be required for somite specification (Nikaido et al., 2002; Ban et al., 2019), and in *R*. sp. ‘chilingali’ and *A. calliptera* it is expressed in a broad domain of gene expression in the anterior PSM, as expected, with a sharp anterior boundary and a more shallow posterior boundary, similar to the pattern that has been observed in zebrafish (Nikaido et al., 2002; Ban et al., 2019) (Fig.5A””, B””, D””, E””). As with other genes patterning the PSM, spatial expression of *tbx6* is indistinguishable between both cichlid species when quantified at early and mid segmentation.

Finally, *MyoD* is involved in muscle development in the newly formed somites (Weinberg et al., 1996; Osborn et al., 2020), and is expressed in nascent somites and in a single row of cells directly adjacent to the notochord - the adaxial cells - in both species (Fig.5.D””’, E””’, F). When quantified, the expression of this gene, like that of all other genes characterised, it too is indistinguishable between the two cichlid species.

Altogether, our data indicate that *A. calliptera* and *R*. sp. ‘chilingali’ do not differ in the molecular patterning of the PSM at early or mid segmentation stages. This supports the notion that the molecular mechanisms underlying the development of the PSM during somitogenesis are conserved in these species.

## DISCUSSION

### Lake Malawi cichlids have evolved increased somitic counts by modifying the initial conditions of somitogenesis

Altogether, our results indicate that *R*. sp. ‘chilingali’ has evolved an increased number of somites by altering the initial conditions of somitogenesis rather than by modifying the process of somitogenesis itself. While *R*. sp. ‘chilingali’ and *A. calliptera* differ in the duration of somitogenesis, which takes seven hours longer in *R*. sp. ‘chilingali’ than in *A. calliptera*, the rate of somitogenesis is conserved between the two (Fig.1.N&O). Furthermore, somites were found to form sequentially throughout somitogenesis in both species, with no evidence of simultaneous formation of somites or a difference in the rate of somitogenesis, even at the onset of somitogenesis (Fig.4).

We have shown that the early *R*. sp. ‘chilingali’ PSM is significantly larger than that of *A. calliptera* in every dimension measured, being 19% longer, 61% greater in volume, and having 57% more cells (Fig.3). Despite these significant initial differences in PSM size, PSM growth rates are indistinguishable between both cichlid species (Fig.3), and the expression patterns of key molecular regulators of PSM morphogenesis are indistinguishable between the two (Fig.5). Nascent somite size is largely comparable (Fig.2.C.) and axial elongation rates are predicted to be similar and not alone able to account for the divergence in vertebral counts. Together, these observations indicate that the dynamics of the molecular clock and the morphogenetic processes underlying somitogenesis are conserved, and that it is only the initial conditions in the form of the size of the PSM at the onset of this process, that have diverged.

If the segmentation clock dynamics or associated morphogenetic mechanisms had diverged, we would expect alterations in PSM anatomy, molecular patterning, and/or growth trajectories; however, none are observed. Even when metrics differ in absolute size (e.g. PSM length, MPZ length), developmental trajectories over time are parallel, reinforcing the interpretation that initial conditions, rather than process, have evolved (Fig.3 anf Fig.S2). Therefore our data suggest that the additional tissue required to generate more somites in *R*. sp. ‘chilingali’ is already present before somitogenesis begins. They support a model in which the extended duration of somitogenesis, and the resulting increase in vertebral number relative to *A. calliptera*, is driven by interspecific differences in PSM size established prior to somitogenesis, rather than by the divergence in any of the mechanisms underlying the process itself.

Importantly, our data show that these interspecific differences are established prior to somitogenesis. *R*. sp. ‘chilingali’ initiates somitogenesis later than *A. calliptera*, suggesting an early heterochronic shift during blastulation or gastrulation (Fig.1). An extended pre-segmentation period could permit increased cell proliferation, resulting in a larger embryonic axis at the onset of somitogenesis. However, the PSM in *R*. sp. ‘chilingali’ is still disproportionately larger than that of *A. calliptera*, failing to scale with embryo size and suggesting a PSM-specific divergence in the mechanisms responsible for the specification of this tissue (Fig.S2). Whole-embryo transcriptomic data for these species further support our interpretations by revealing that the largest inter-specific differences occur at the four-somite stage, near the onset of segmentation (Marconi et al., 2024), becoming more similar thereafter. Together, these findings indicate that divergence between these species is most pronounced at the onset of somitogenesis, supporting the view that somitic counts have evolved by evolving the initial conditions of somitogenesis and suggesting that evolutionary differences in other phenotypic traits may likewise be established during the earliest stages of embryonic development.

### Evolution can be driven by changes in the initial conditions of developmental pro-grams

Our findings suggest a simple mechanism to evolve traits. Phenotypic evolution need not be rooted in the modification of the underlying developmental process itself, but rather in changes to its initial conditions. Instead of modifying a complex developmental process, such as somitogenesis and axial elongation, it may be simpler to instead evolve the initial conditions of that process, particularly over shorter timescales. These potentially subtle changes early on in embryogenesis can cascade through development giving rise to dramatic consequences for the adult phenotype. In the context of the evolution of vertebral counts in Lake Malawi cichlids we have shown that these initial conditions correspond to the initial size of the body axis, and of the PSM in particular.

Other species have also been found to achieve phenotypic innovations by modifying development at early stages prior to the onset of segmentation. The two-tailed goldfish has evolved a second tail by altering dorsoventral patterning at the gastrula and tailbud stages, driven by a mutation to the gene chordin which arose during artificial selection (Abe et al., 2014). This morph exhibits delayed blastopore closure relative to single-tailed goldfish, and has a larger PSM, which interestingly has resulted in the evolution of two tails rather than increased vertebral counts (Abe et al., 2014; Li et al., 2019).

Heterochronies during gastrulation in the Mexican cavefish, *Astyanax mexicanus*, have been found to generate organizer, and interestingly, axial mesodermal tissues with diverging gene expression patterns (Torres-Paz et al., 2019). These differences have been postulated to have contributed towards the divergence in brain evolution between cave and surface morphs. In that work, authors founds that these differences depended fully on ma-ternal factors, and that certain traits such as retinal morphogenesis, retained significant maternal influence throughout development and well into the larval stages. Perhaps in the closely related Lake Malawi cichlids, maternal factors also play an important role affecting early stages in embryogenesis and cascading on during development affecting certain processes and manifesting in adult traits.

The end of somitogenesis corresponds to the phylotypic stage in vertebrates, also known as the pharyngula stage. This is the stage when vertebrate embryos across species most resemble each other both morphologically (Baer, 1828) and transcriptionally (Irie & Kuratani, 2011). These striking similarities are believed to reflect the developmental stages of most constraint, which may be underpinned by the obligate and deeply conserved progressive expression of the hox genes (Duboule, 1994). Given this deep evolutionary conservation and strong constraints on the morphology of the embryo after segmentation, perhaps the easiest way to evolve vertebral counts over short evolutionary timescales might require by-passing any changes to somitogenesis and any other processes that might take place close to the phylotypic stage, in favor of changes to the initial conditions of these processes.

Finally, changing initial conditions might not be the only way to evolve a phenotype without modifying the underlying developmental process that generates it. Boundary conditions, which dictate how a system interacts with its spatial or structural environment, might also evolve and affect the outcome of conserved developmental processes. It has been proposed that changes in tissue tectonics - or how tissues develop relative to each other - might be enough to vary the placement of signalling centres and responding tissues, potentially resulting in changes to the coordinated timing of development (Busby & Steventon, 2021) that could lead to phenotypic evolution. The effect of boundary conditions is well illustrated in two-dimensional stem cell colonies, where the geometry of the micro-patterns used has been seen to modify the position of primitive streak-like regions, as well as the direction and symmetry of cell migration (Warmflash et al., 2014; Etoc et al., 2016; Heemskerk et al., 2019) Furthermore, it has been shown that altering geometry and boundary conditions in embryos by modulating the spatial distribution of forces, flows, and active processes - such as through changes in the position of cell ingression - can shift between the different modes of gastrulation described across vertebrates (Serrano Nájera et al., 2025; Serrano Nájera & Weijer, 2023; Chuai et al., 2023; Serrano Nájera & Weijer, 2023). Finally, heterochrony in expression of “timer genes” controlling how arthropod segmentation GRN(s) are deployed within the embryo is also thought to account for the variation in segmentation methods in arthropods (Clark et al., 2019; Clark & Peel, 2018; Taylor & Dearden, 2022), another example of how boundary or external conditions can shape developmental processes. Taken together, this suggests that boundary conditions, alongside initial conditions as demonstrated in this study, may represent an important and currently underappreciated driver of phenotypic evolution.

## CONCLUSION

We have shown that *R*. sp. ‘chilingali’ have likely evolved increased somitic counts by modifying the initial conditions of somitogenesis, rather than the process of somitogenesis itself. *R*. sp. ‘chilingali’ have a significantly larger PSM than *A. calliptera* at the onset of somitogenesis, providing the excess tissue used to produce more somites. Moreover, there is no evidence for a change in the process of somitogenesis between species, as morpho-metric analyses of the anatomy of the PSM, and gene expression patterning of the PSM have revealed that these are very similar between *R*. sp. ‘chilingali’ and *A. calliptera*. This work therefore presents an alternative mechanism by which phenotypic evolution can be achieved, showing that altering the initial conditions of development can result in dramatic phenotypic changes, without the need to modify the underlying developmental process. This mechanism might be particularly useful over short evolutionary timescales or when a developmental process is highly constrained, as could be the case with somitogenesis. We hope our work will encourage others to look for alternative developmental drivers of evolutionary change in closer related species to reveal how phenotypic evolution first emerges.

## METHODS

### Animal husbandry

*A. calliptera* and *R*. sp. ‘chilingali’ were reared at 26*^◦^*C, under a 12:12 hour day-night cycle. *A. calliptera* were fed JBL Novo Malawi flake (for algae-eating cichlids), and *R*. sp. ‘chilingali’ were reared on JBL Novo Malawi flake for carnivores. Feed was supplemented with pellets, frozen brine shrimp, and frozen fish for enrichment and to encourage breeding.

### Extraction and fixation of cichlid embryos

Embryos were extracted from stock tanks at least weekly. Embryos were removed from mouthbrooding females by first catching the female, and then flushing the mouth with water using a pasteur pipette until all embryos were released. All animal work was conducted following approval by the Departmental Animal Welfare Ethical Review Body (AWERB) for the Department of Biology at the University of Oxford.

To establish the rate of somitogenesis, embryos were staged in hours post fertilisation (hpf) by monitoring breeding tanks hourly for the appearance of new mouthbrooders, or if a clutch of one-cell embryos was extracted during regular fish work, the time of extraction was treated as the estimated time of fertilisation. If breeding was observed then fish were left for the duration of courtship (*≈* 1.5 hrs) and then embryos were extracted, with the end of courtship treated as the time of fertilization. As the first division takes *≈* 1*−*2 hrs (Fujimura & Okada, 2007; Marconi et al., 2023), these collection methods are equivalent.

If embryos were to be reared to a specific stage, they were incubated in a solution of tank water with 1mg/L of methylene blue (Sigma-Aldrich) to prevent mould and bacterial contamination. Embryos were incubated on a slow rocker at 26*^◦^*C. Water was changed and dead embryos removed daily.

Fixation was performed overnight at 4*^◦^*C, in 4% v/v formaldehyde (Sigma).

### Hybridization chain reaction

Hybridization chain reaction (HCR) was performed as per (Choi et al., 2016, 2018; Fulton et al., 2022b). Probes were designed by Molecular Instruments; accession numbers for genes can be found in Table 1. In short, embryos were fixed overnight at 4*^◦^*C in 4% formaldehyde and then dehydrated in a 25%, 50%, 75% methanol:Phosphate Buffered Saline (PBS) series, leaving tissue to equilibrate for five minutes for each methanol concentration. Embryos were then stored at *−*20*^◦^*C in 100% methanol. Note that all washes performed are for five minutes unless indicated otherwise.

**Table 1:** Genes used in this study.

| Gene name | NCBI accession |
| --- | --- |
| *tbxta* | XM_026157450 |
| *tbx16* | XM_026187037 |
| *tbx6* | XM_026153614 |
| *sox2* | XM_026147476.1 |
| *myoD* | XM_026172524.1 |
| *msgn* | XM_026142939 |

To perform the HCR, embryos were first rehydrated in a 75%, 50%, 25% methanol:PBS series as before. Embryos were then dissected from the yolk in PBS, and placed in a glass staining dish. Permeabilization with proteinase K was typically not required for these stages. If permeabilization was required, embryos were permeabilized for one hour in 10*µ*g/mL proteinase K (Sigma) before being washed twice in PTw and re-fixed in 4% formaldehyde for 30 minutes.

Embryos were pre-hybridized for 30 minutes at 37*^◦^*C in 30% hybridization buffer (Molecular Instruments), before 1*µ*L of probe was added per 100*µ*L of hybridization buffer. Note that to ensure strong and consistent staining we used 100*µ*L of hybridization buffer per half-dozen embryos, and did not stain more than 15 embryos per staining dish. The embryos were then left overnight to hybridize at 37*^◦^*C.

The following day, we washed out the probe using 4×15 minute washes in probe wash buffer (Molecular Instruments) at 37*^◦^*C. At room temperature, we performed 3×5 minute washes in SCT (5X SSC + 0.1% Tween-20), before pre-amplifying in amplification buffer (Molecular Instruments). We heat-shocked the hairpins at 95*^◦^*C for 90 seconds in separate PCR tubes, before adding these to the embryos. Embryos were then incubated overnight, in the dark.

The following morning we washed out the hairpins in SSCT and added 1*µ*L/mL DAPI at least overnight, before mounting in 90% glycerol and imaging.

### Phalloidin staining

Embryos were extracted and fixed as above, before being washed three times in PTw and dissected. Embryos were then permeabilized for a total of two days in 1% PTx, washing four times over this time period. Embryos were then washed three times in PTw before being stained overnight in half a unit (10*µ*L) Phalloidin-CF568 (Biotin, BT00044-T) in 100*µ*L PTw and 0.5*µ*L DAPI (Biotin, BT40011). Finally, the stain was washed out three times in PTw, before embryos were mounted in 90% glycerol.

### Imaging

Imaging was performed on either an Olympus FV3000, Leica LSM980, or Leica LSM880. Microscope configuration is summarized in Table 2

**Table 2:**
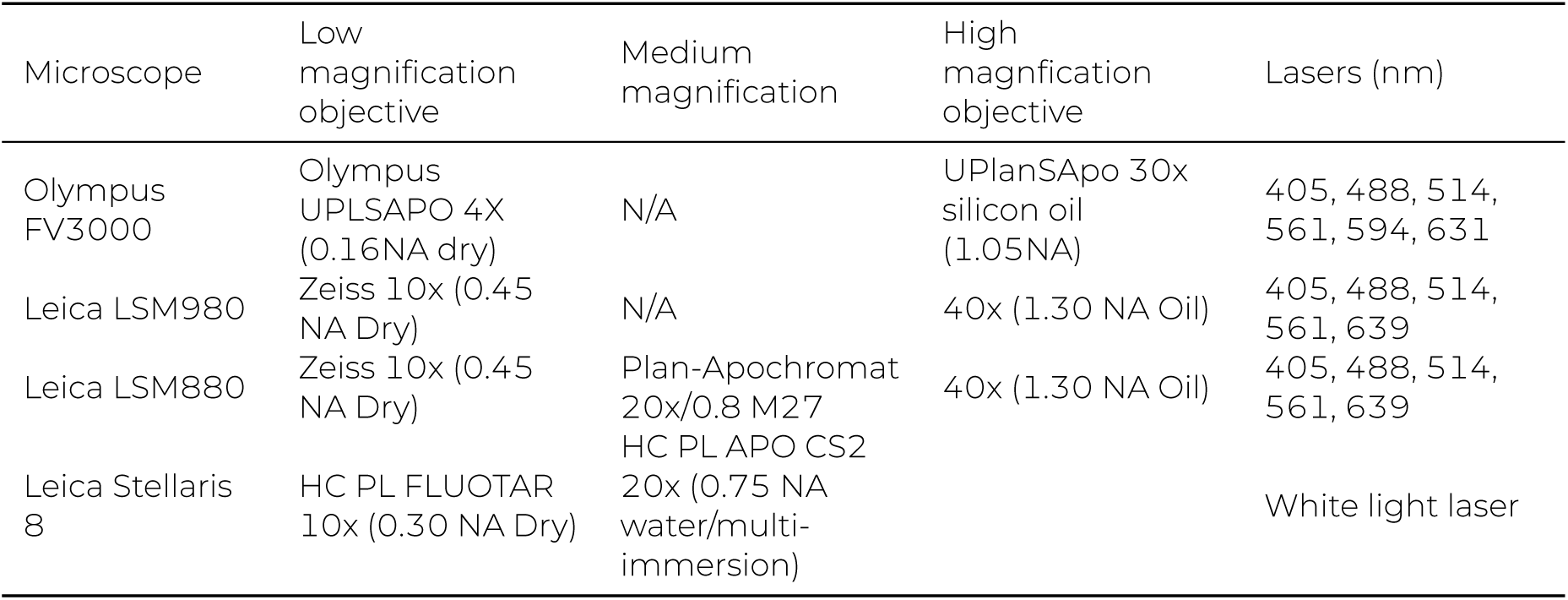
Microscope configuration.

### Morphometrics

Morpometric measurements were performed by manually annotating images within OMERO (Burel et al., 2015). Anatomical regions were defined as follows: The PSM length is defined as the distance between the posterior-most tip of the PSM, at the level of the notochord, and the posterior boundary of the nascent somite; measured through the notochord. The notochord length is the distance between the posterior boundary of the nascent somite, and the posterior tip of the notochord, measured through the notochord. The MPZ length is the difference between these two values. Somite length is defined as the distance between the posterior and anterior boundaries of the nascent somite, measured at the middle of the somite. PSM width was measured by placing two points at the intersection of the posterior boundary of the nascent somites, and the lateral aspect of the PSM, and by computing the euclidean distance between these points in Python. PSM depth is similarly measured, by placing points at the dorsal and ventral most points of the PSM, at the anterior-posterior position of the lateral tip of the notochord. Notochord width is measured at the middle of the notochord. Metrics are shown in Fig.3.

The embryo length was measured as a poly-line across the AP axis of the embryo, from the tip of the nose to the tip of the tailbud, following the embryonic spinal cord.

Principal component analysis was performed in R v4.4.2 using the ‘prcomp‘ command from the ‘stats‘ package, scaling data to have unit variance prior to the PCA being performed.

### Manual tissue segmentation

Tissues were manually segmented using the Segmentation Editor tool in ImageJ (Schindelin et al., 2012). The whole tailbud was first outlined every 5-10 slices, then the labels were interpolated and the label added to the image. The notochord, neural tube, and somites were then segmented using the same method, and given unique labels for identification.

To measure the volume of different tissues, the label object was resized so that each voxel measured 1*µ*m^3^. The number of voxels in each label were then counted. For somites, the number of somites segmented (one or two) was detected automatically by extracting the somite label, labeling each somite, and then counting the number of labels. The mean somite volume was then computed by dividing the total somite volume by the number of somites.

### Measuring spatial molecular patterns across the AP axis

Gene expression patterns across the anterior-posterior axis were measured in Fiji as follows. First, an average intensity projection of relevant focal planes was performed in ImageJ. Average intensity projections were always performed on focal planes adjacent to the notochord. A polyline was then drawn across the PSM, from the posterior boundary to the nascent somite to the posterior of the notochord, following the middle of the PSM. Expression intensity was then measured, averaging across 20-100 pixels adjacent to the line. The number of pixels was chosen such that the polyline width was as close as possible to the width of the PSM itself, without including any tissue outside the PSM.

### Cell counting

Cell counting was performed using a method derived from a standard spot counting protocol by Robert Haase, using the pyclesperanto library (Haase et al., 2020). The DAPI (nu-clear) channel was first extracted from the image, and rescaled to have voxels of size 0.5*µ*m^3^. A gaussian blur was then performed on the nuclear channel, with a radius of 0.5*µ*m (one voxel) in the X and Y dimensions, and zero voxels in the Z dimension. The local maxima was detected, using a ‘box’ (cubed) connectivity, and radius of 1.5*µ*m (three voxels) in all dimensions. To merge nearby spots, a binary dilation was performed with radius 1.5*µ*m (three voxels) in the X and Y dimensions and 0.5*µ*m (one voxel) in the Z dimensions, before spots were labelled, the relevant tissue extracted, and the number of unique labels (e.g. number of cells) in the tissue was counted.

The quality of spot detection was verified blind after segmentation, and embryos where spot detection was not accurate were discarded.

## ACKNOWLEDGMENTS

We are very grateful to Christine Soper, Allex Turner, and Helen Saunders for excellent fish care, and to members of the Verd lab for insightful comments during the research process. The authors gratefully acknowledge the use of the University of Oxford Advanced Research Computing (ARC) facility (http://dx.doi.org/10.5281/zenodo.22558), and the Micron Advanced Bioimaging Facility (supported by Wellcome Strategic Awards 091911/B/10/Z and 107457/Z/15/Z) for their support and assistance in this work. SET was supported by a Clarendon Scholarship, the William Georgetti Scholarship, and a Raymond-Sisam Graduate Scholarship from St Hildas College. JH was supported by a Natural Environment Research Council DTP studentship (grant NE/S007474/1). BV was supported by the John Fell Fund, University of Oxford (0009780), the Royal Society (RGS111324) and the European Research Council (101163722 StG Counts).

## AUTHOR CONTRIBUTIONS

Conceptualization: BV. Methodology: SET, JH, BV. Investigation: SET, JH. Visualization: SET. Writing: SET, JH, BV. Editing: SET, JH, BV. Funding Acquisition: SET, JH, BV. Supervision: BV.

## AUTHOR COMPETING INTERESTS

No competing interests.

## SUPPLEMENTARY MATERIAL

**Table S1:** Statistics on PSM metrics.

|  | variable stage |  |  |  | comparison |  |  | variable stage + species |  |  |  |  |  |  |
| --- | --- | --- | --- | --- | --- | --- | --- | --- | --- | --- | --- | --- | --- | --- |
|  | R<sup>2</sup> | slope | 95 CI | F statistic | p-value | RSS model 1 | RSS model 2 | R<sup>2</sup> | slope | 95 CI | species difference | 95 CI species diff. | p count | p species |
| PSM length | 0.71 | -13.02 | -14.53 — -11.51 | 285.85 | 4.4e-33 | 570125 | 394490 | 0.80 | -13.68 | -14.95 — -12.4 | 77.95 | 56.6 — 99.3 | 1.1e-39 | 8.4e-11 |
| notocord length | 0.76 | -10.13 | -11.17 — -9.1 | 370.54 | 6.5e-38 | 266211 | 211735 | 0.81 | -10.50 | -11.43 — -9.56 | 43.41 | 27.77 — 59.05 | 4.4e-42 | 3.1e-07 |
| PSM width | 0.43 | -3.43 | -4.15 — -2.72 | 87.78 | 7.2e-16 | 129073 | 123839 | 0.45 | -3.55 | -4.26 — -2.83 | 13.45 | 1.49 — 25.42 | 3.3e-16 | 0.03 |
| PSM depth | 0.12 | 1.11 | 0.55 — 1.67 | 15.12 | 1.7e-04 | 78637 | 77451 | 0.13 | 1.06 | 0.49 — 1.62 | 6.41 | -3.05 — 15.87 | 1.6e-04 | n.s. |
| MPZ length | 0.34 | -2.89 | -3.63 — -2.15 | 58.49 | 6.5e-12 | 137317 | 102837 | 0.50 | -3.18 | -3.83 — -2.53 | 34.54 | 23.63 — 45.44 | 1.6e-14 | 8.7e-09 |
| Somite length | 0.48 | -0.75 | -0.89 — -0.61 | 108.45 | 2.5e-18 | 5022 | 4797 | 0.51 | -0.73 | -0.87 — -0.59 | -2.78 | -5.14 — -0.43 | 9.5e-19 | 0.02 |
| notocord width | 0.30 | -0.54 | -0.69 — -0.39 | 50.77 | 9.4e-11 | 5560 | 5503 | 0.31 | -0.55 | -0.7 — -0.4 | 1.41 | -1.12 — 3.93 | 9.4e-11 | n.s. |
| length:width | 0.16 | -0.03 | -0.04 — -0.01 | 21.73 | 8.4e-06 | 29 | 27 | 0.20 | -0.03 | -0.04 — -0.02 | 0.23 | 0.05 — 0.4 | 5.6e-06 | 0.01 |
| length:depth | 0.37 | -0.35 | -0.43 — -0.26 | 68.39 | 2.5e-13 | 1680 | 1680 | 0.37 | -0.35 | -0.43 — -0.26 | 0.14 | -1.25 — 1.54 | 3.2e-13 | n.s. |
| depth:width | 0.23 | -0.11 | -0.15 — -0.07 | 33.69 | 5.7e-08 | 356 | 354 | 0.23 | -0.11 | -0.15 — -0.07 | -0.25 | -0.89 — 0.39 | 6.1e-08 | n.s. |

**Figure S1:**
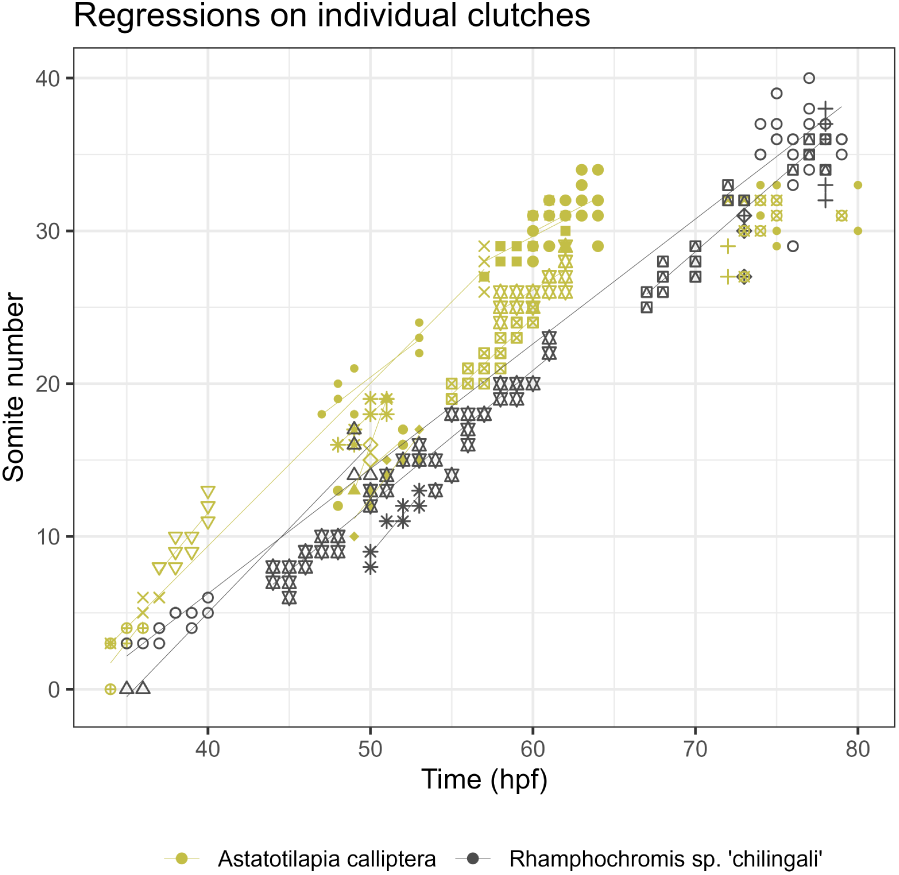
Number of somites versus time in both species. The linear regression was constructed for both species, for each clutch individually, using ordinary least squares and the equation is displayed at the bottom right. Yellow: *A. calliptera*, black: *R*. sp. ‘chilingali’.

**Table S2:** Probability of interaction between species and somite stage.

|  | R<sup>2</sup> | p species | p count | p interaction |
| --- | --- | --- | --- | --- |
| PSM length | 0.80 | 1.0e-10 | 2.3e-39 | n.s. |
| notocord length | 0.81 | 3.4e-07 | 1.0e-41 | n.s. |
| MPZ length | 0.50 | 9.9e-09 | 2.0e-14 | n.s. |

**Figure S2:**
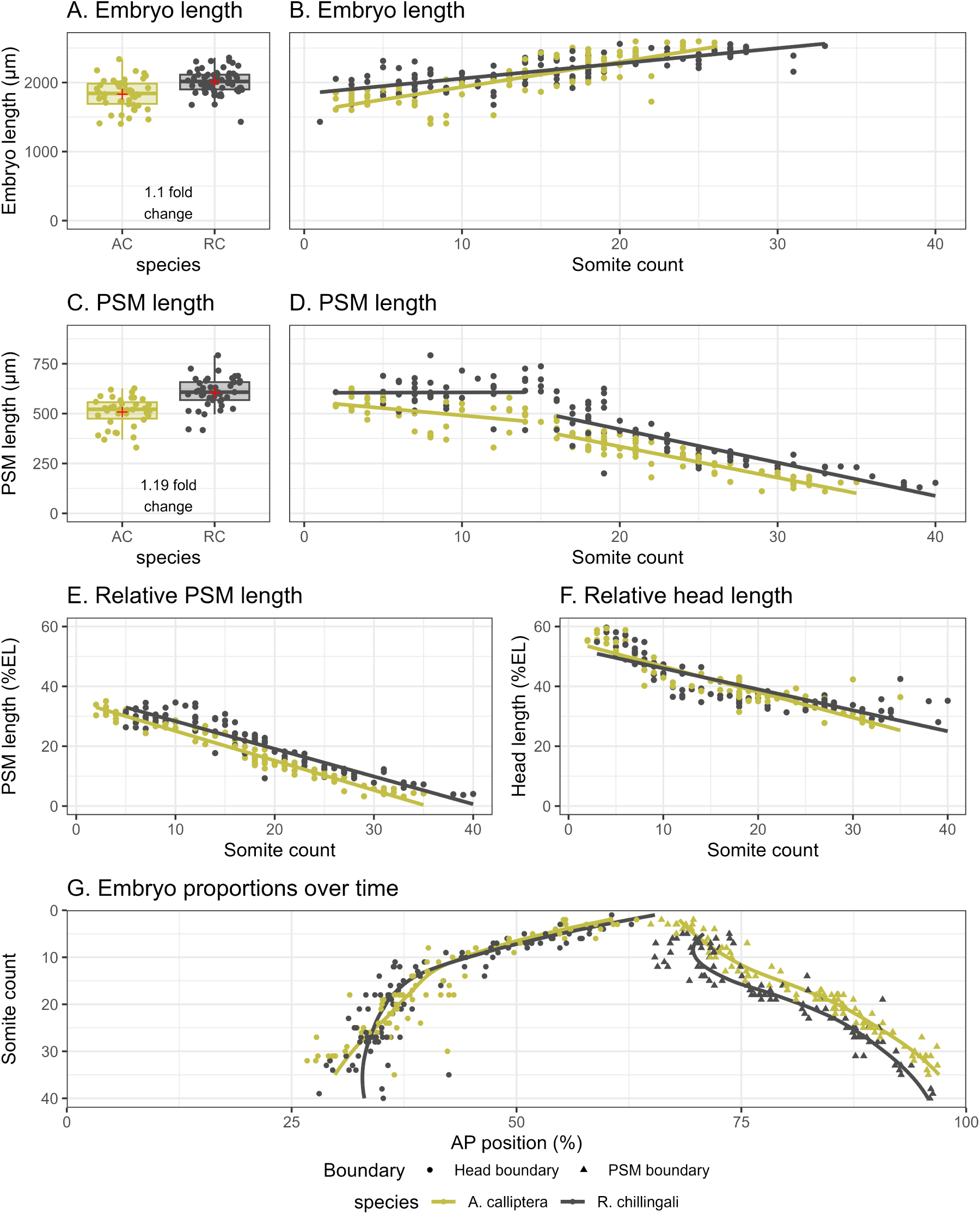
Embryo dimensions differ between species. **A.** Embryo length, in *µ*m, for both species, for embryos less than 15ss. **B**. Embryo length, in *µ*m, for both species, at different somite stages. C. PSM length, in *µ*m, for both species, for embryos less than 15ss. **D**. PSM length, in *µ*m, for both species, at different somite stages. **E**. PSM length against somite stage as a percentage of embryo length. **F**. Head length against somite stage as a percentage of embryo length. **G**: Relative head and PSM proportions of embryonic length over time (in somite stage). For **C, D, E, F** the lines represent the linear regression of the variable for each species. For **G**, the coloured lines represent the local regression-based smoothing for each species.

**Figure S3:**
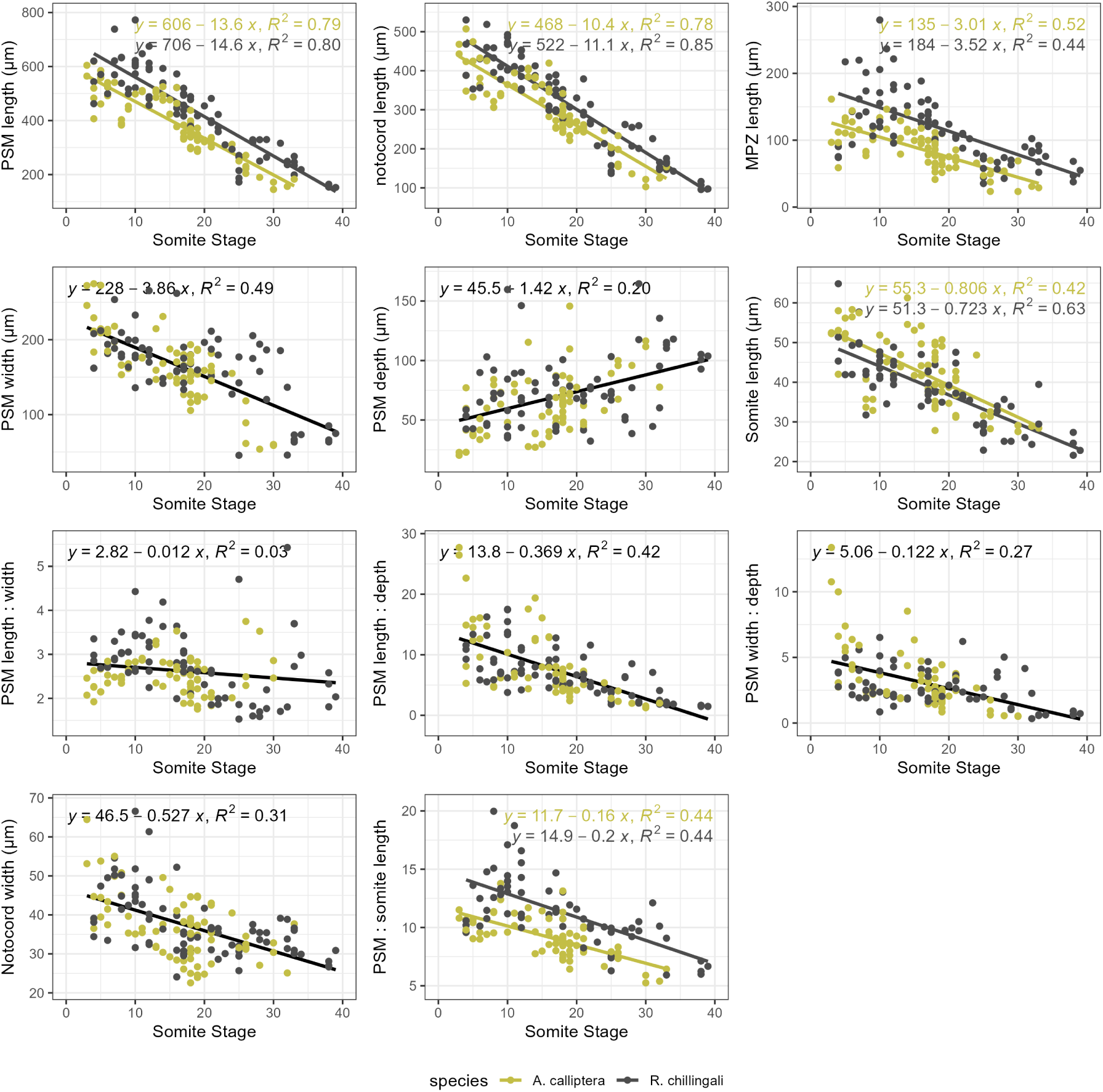
PSM measurements used for PCA. Solid black lines indicate the trendline for measurements where species did not statistically differ, and different coloured lines indicate species-specific trendlines. *A. calliptera*: yellow. *R*. sp. ‘chilingali’: grey.

**Figure S4:**
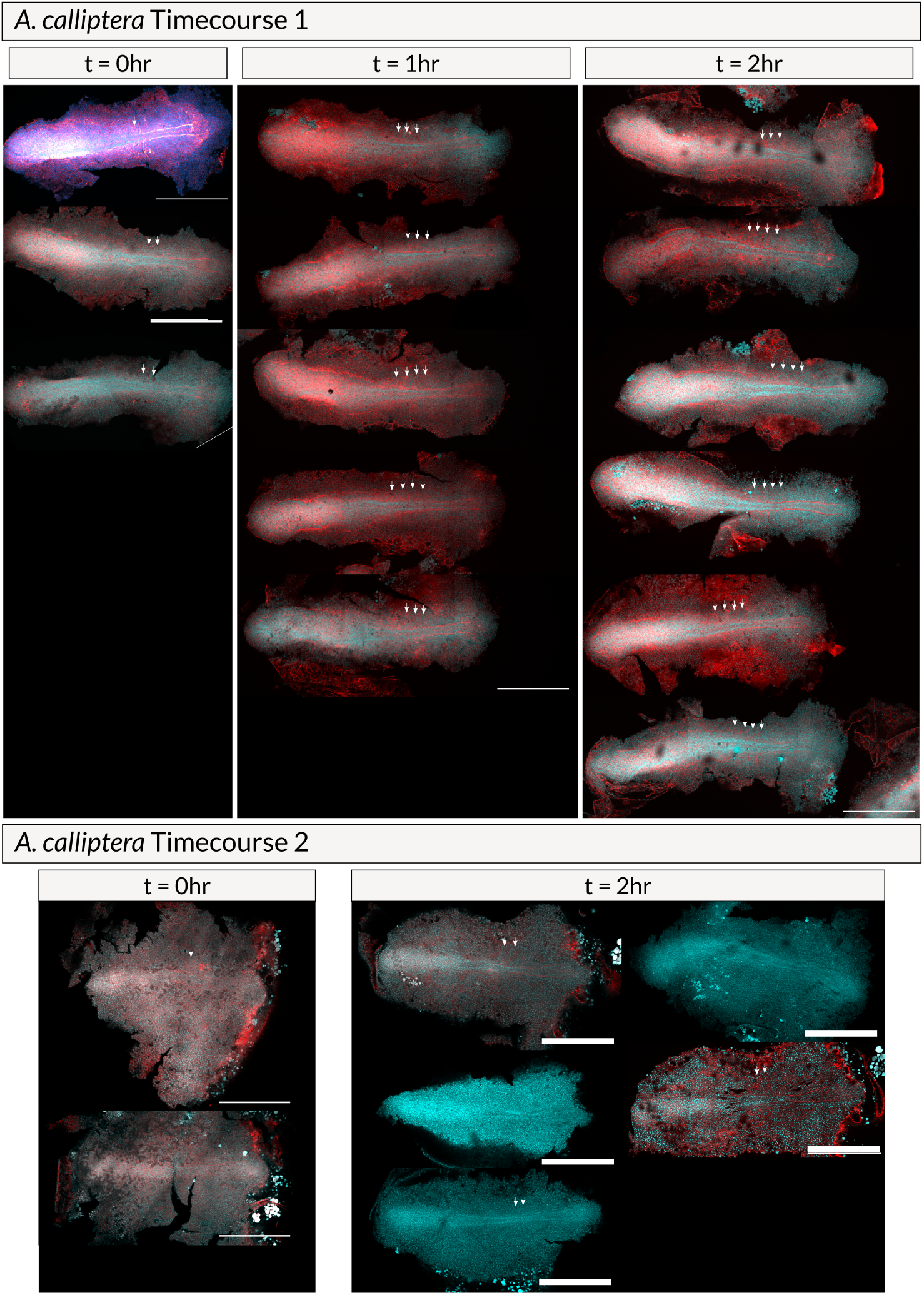
*A. calliptera* somite numbers at the onset of segmentation. Embryos were fixed at hourly intervals, before being dissected and stained with DAPI (cyan) and phalloidin (red). Somites are indicated with arrows. Scale bar: 500 *µ*m. All embryos are oriented anterior left.

**Figure S5:**
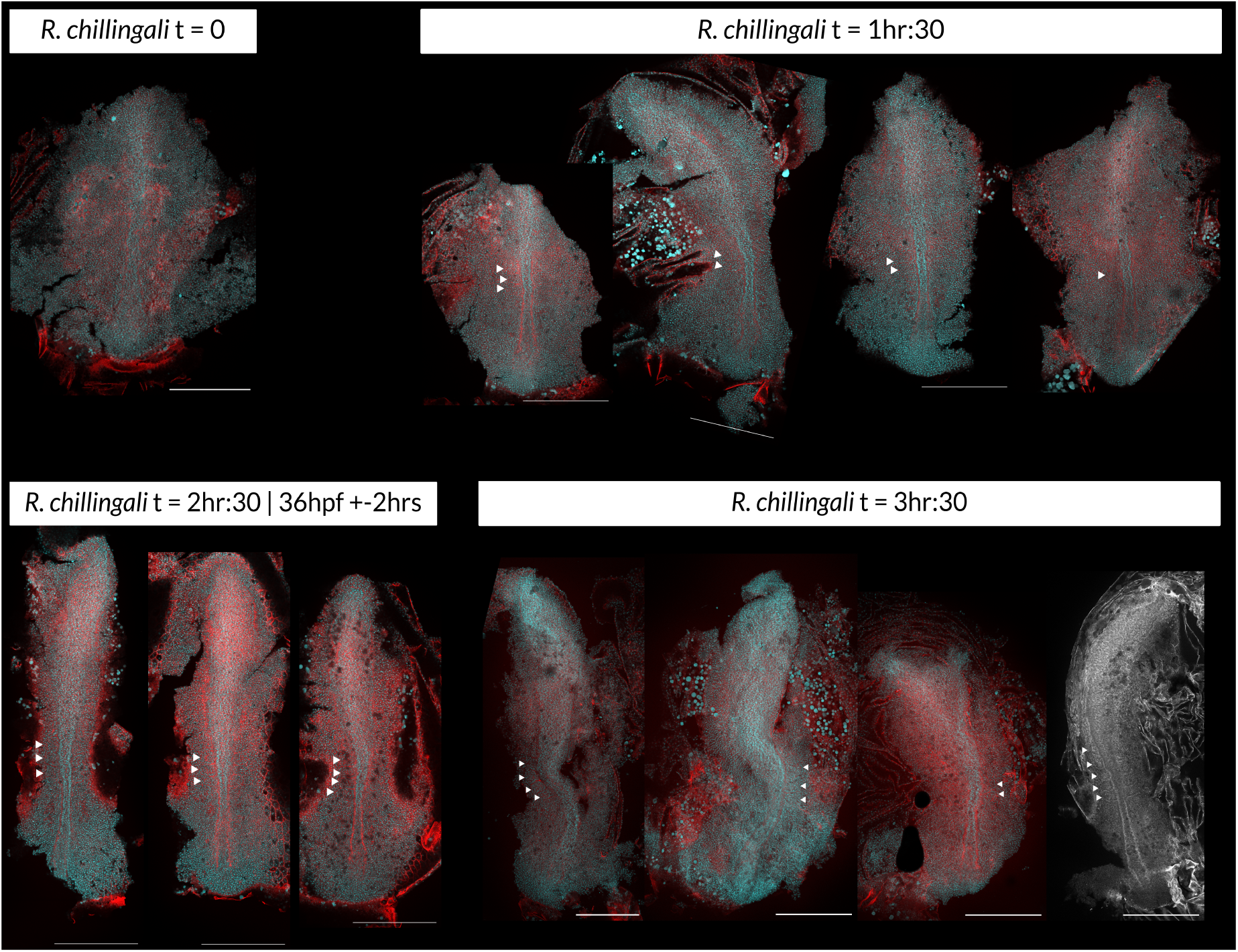
*R*. sp. ‘chilingali’ somite numbers at the onset of segmentation. Embryos were fixed at hourly intervals, before being dissected and stained with DAPI (cyan) and phalloidin (red). Somites are indicated with arrows. Scale bar: 500 *µ*m. All embryos are oriented anterior towards the top.

